# De novo enteric neurogenesis in post-embryonic zebrafish from Schwann cell precursors rather than resident cell types

**DOI:** 10.1101/2020.06.01.127712

**Authors:** Wael Noor El-Nachef, Marianne E. Bronner

**Affiliations:** Vatche and Tamar Manoukian Division of Digestive Diseases, University of California Los Angeles, Los Angeles, CA 90095 USA; Division of Biology and Biological Engineering, California Institute of Technology, Pasadena, CA 91125 USA

**Keywords:** enteric nervous system, neural crest, prucalopride, 5-HT_4_

## Abstract

The enteric nervous system is essential for normal gastrointestinal function, but evidence regarding postnatal enteric neurogenesis is conflicting. Using zebrafish as a model, we explored the origin of enteric neurons that arise in post-embryonic life in normal development and injury, and tested effects of the 5-HT_4_ receptor agonist, prucalopride.

To assess enteric neurogenesis, all enteric neurons were photoconverted prior to time-lapse imaging to detect emergence of new neurons. Injury was modeled by two-photon laser ablation of enteric neurons. Lineage tracing was performed with neural tube injections of lipophilic dye and with an inducible Sox10-Cre line. Lastly, we tested prucalopride’s effect on post-embryonic enteric neurogenesis.

The post-embryonic zebrafish intestine appears to lack resident neurogenic precursors and enteric glia. However, enteric neurogenesis persists post-embryonically during development and after injury. New enteric neurons arise from trunk neural crest-derived Schwann cell precursors. Prucalopride increases enteric neurogenesis in normal development and after injury if exposure occurs prior to injury.

Enteric neurogenesis persists in the post-embryonic period in both normal development and injury, appears to arise from gut-extrinsic Schwann cell precursors, and is promoted by prucalopride.

**SUMMARY STATEMENT:** Trunk crest-derived enteric neurogenesis is poorly understood. We find post-embryonic zebrafish lack resident neuronal precursors yet enteric neurogenesis from trunk crest-derived precursors occurs in development, injury, and is promoted by prucalopride.

## INTRODUCTION

The enteric nervous system (ENS) is composed of as many neurons as the spinal cord and is responsible for mediating crucial functions of the gastrointestinal tract, including motility, afferent “sensing”, and secretion (Furness, 2006). Pathologies involving the ENS range from congenital neurocristopathies such as Hirschsprung disease (Lake and Heuckeroth, 2013) to acquired conditions such as esophageal achalasia (Kraichely and Farrugia, 2006) and diabetic gastroparesis (Farrugia, 2015). Establishing fundamental features of ENS homeostasis and the potential for enteric neuronal regeneration could assist in the discovery of novel therapies to treat enteric neuropathies. As the intestine lengthens during postnatal life (Struijs et al., 2009; Weaver et al., 1991) and is susceptible to injury during episodes of inflammation (Brierley and Linden, 2014) and mechanical stress (Wood, 2011), there is likely to be a need for continuous ENS neurogenesis throughout life to increase numbers and/or replace lost enteric neurons. Surprisingly, the question of postnatal enteric neurogenesis remains controversial with conflicting reports arguing that postnatal enteric neurogenesis does not occur (Joseph et al., 2011), occurs after injury (Goto et al., 2013; Katsui et al., 2009; Laranjeira et al., 2011), occurs after exposure to 5-HT_4_ receptor agonists (Goto et al., 2013; Katsui et al., 2009; Liu et al., 2009; Matsuyoshi et al., 2010), or occurs rapidly with a turnover measured in days (Kulkarni et al., 2017).

During development, the ENS is classically described as arising from the vagal neural crest which emerges from the caudal hindbrain, migrates to and invades the foregut and then migrates along the rostrocaudal extent of the gut to colonize the entire length of the intestinal tract (Lake and Heuckeroth, 2013). In addition to the vagal neural crest, there is a modest contribution from the sacral neural crest to the hindgut in some species (Burns et al., 2000; Wang et al., 2011). During initial colonization of the intestine, studies in mice have revealed that the spatial and functional organization of the mammalian ENS occurs via clonal expansion of precursors of neuronal and/or glial character, and depends on factors such as isometric growth and lineally unrelated neighboring cells (Lasrado et al., 2017).

More recently, evidence has arisen for a novel source of enteric neurons originating from neural crest stem cells that remain nascent along peripheral nerves and are often referred to as “Schwann cell precursors (SCPs)” (Furlan and Adameyko, 2018). These progenitors reside within and migrate along peripheral nerves to give rise to diverse cell types, including parasympathetic neurons (Dyachuk et al., 2014; Espinosa-Medina et al., 2014), melanocytes (Adameyko et al., 2009), and cardiomyocytes (Tang et al., 2019). SCPs account for 20% of neurons in the colon of mice (Uesaka et al., 2015), approximately half of foregut neurons in chick (Espinosa-Medina et al., 2017), and all enteric neurons in lamprey (Green et al., 2017), a basal vertebrate that lacks a vagal neural crest. Moreover, these SCPs often arise from trunk rather than vagal levels. This suggests that the original evolutionary strategy to populate the intestine with neurons is via SCPs rather than vagal neural crest cells, and a trunk neural crest contribution has been retained in avian and mammalian species.

While best studied in chick and mouse, development of the enteric nervous system is largely conserved across jawed vertebrates including teleosts like zebrafish (Ganz, 2018; Heanue et al., 2016). As in chick and mouse, zebrafish vagal neural crest cells enter the foregut and migrate rostrocaudally to colonize its entire length. Zebrafish offer several advantages for studying ENS development and maturation. First, they are amenable to live-imaging techniques that allow direct visualization *in vivo* of cell behavior within the context of the entire organism. Second, zebrafish have a simplified ENS compared with that of chick and mouse, with two streams of vagal neural crest cells migrating along the left and right sides of the intestine, facilitating imaging studies. Third, a variety of transgenic lines are available to label particular cell types of interest. Finally, zebrafish are amenable to experimental manipulation and highly accessible to drug treatment.

Here, we take advantage of the ease of imaging and manipulation of the zebrafish model to examine the role of de novo neurogenesis as the ENS transitions from embryonic to larval stages. As expected, we find that new neurons are added as the animal grows as well as after injury. Surprisingly, we show that these new neurons do not arise from resident progenitors or enteric glia and indeed that zebrafish appear to lack these cell populations. Rather than originating from enteric precursors in the intestine, we provide evidence that post-embryonic enteric neurons arise from trunk neural crest-derived Schwann cell precursors that migrate into the intestine and differentiate into new neurons. Lastly, we find that prucalopride, a 5-HT_4_ receptor (5HT4R) agonist recently approved for use in the United States(“Drug Approval Package,” n.d.), promotes enteric neurogenesis and is protective in an injury model.

Taken together, our results reveal novel roles for Schwann cell precursors in the context of ongoing neurogenesis in the post-embryonic intestine in both normal development and after injury, and suggest that enteric glia evolved after the teleost lineage on the vertebrate tree, perhaps as vertebrates moved onto land.

## MATERIALS AND METHODS

### Transgenic lines

Zebrafish *(danio rerio)* were maintained at 28°C, with adults on a 13-hour light/11-hour dark cycle. All zebrafish work was completed in compliance with the California Institute of Technology Institutional Animal Care and Use Committee. Transgenic lines used in this study were the photoconvertible Phox2b kaede line (Harrison et al., 2014), the Sox10:GAL4-UAS-Cre (Cavanaugh et al., 2015) (“indelible Sox10 Cre”) line which was crossed with the ubi:switch reporter line (Cavanaugh et al., 2015), the cmlc:GFP Sox10:ERT2-Cre (Mongera et al., 2013) (“inducible Sox10-Cre”) line which was crossed with the ubi: switch reporter line, the Sox10-mRFP line (Kucenas et al., 2008), and the HuC:GCaMP6 transgenic line (Freeman et al., 2014). All lines were within an ABWT background, with the exception of the HuC:GCaMP6 line, which was backcrossed onto the pigmentless “casper” line (White et al., 2008).

### In situ hybridization (ISH) and Immunohistochemistry (IHC)

Embryos and larvae underwent hybridization as previously described (Jowett and Lettice, 1994), with the following changes: samples were stored in ethanol and digestion was performed with 1 mg/mL collagenase 1a [Sigma C9891] (5 min and 12 min for 2 dpf and 3.5 dpf, respectively) prior to proteinase K digestion (12 min and 14 min for 2 dpf and 3.5 dpf, respectively). All imaging of ISH specimens was performed on a Zeiss Imager.M2 with an ApoTome.2 module.

Our whole-mount IHC staining of embryos and larvae protocol was adapted from a prior study (Ungos et al., 2003) and was performed by fixation in 4% PFA in PB overnight at 4°C, then washing in 1x PBS, followed by incubation in 0.5x PBS for 30 minutes. Samples were then placed in blocking solution (2% goat serum, 1% BSA, 1% DMSO, 0.1% Triton X-100, and 0.05% Tween in 1x PBS) for two hours at room temperature. Samples were then incubated in primary antibody diluted in blocking solution overnight at room temperature and washed for 2-3 hours in 1x PBS plus 0.1% Triton X-100. Then, samples were incubated overnight in secondary antibody diluted in blocking solution plus DAPI [1:1000; ThermoFisher Scientific D1306] overnight at room temperature and washed for 2-3 hours in 1x PBS plus 0.1% Triton X100. Samples were then mounted in RIMS (Yang et al., 2014) to achieve optical clearing.

For histologic sections, cryosections were collected at 10 μm thickness. Blocking and antibody incubation occurred the same as with wholemount samples, except that antibody incubations occurred at 4°C and samples were mounted with Fluormount-G [ThermoFisher Scientific, 00-4958-02]. The primary antibodies used were mouse anti-HuC/D IgG2b [1:200; ThermoFisher Scientific A21271], mouse anti-mCherry IgG1 [1:200; Clontech Living Colors 632543], rabbit anti-GFAP IgG [1:200; Genetex GTX 128741], The secondary antibodies used in this study were goat anti-mouse IgG2b 647 [1:500; ThermoFisher Scientific A21242], goat anti-mouse IgG1 568 [1:500. ThermoFisher Scientific A21124], goat anti-rabbit IgG 647 [1:500; ThermoFisher Scientific A21134]). All imaging of IHC specimens was performed on the Zeiss LSM 800 confocal microscope and figures produced with ImageJ software [National Institutes of Health].

### Adult intestine wholemount imaging

Adult zebrafish intestine was procured as previously described (Gupta and Mullins, 2010). Intestine was then opened longitudinally, fixed in 4% PFA in PB overnight at 4°C, washed in 1x PBS, incubated in DAPI 1:1000 for 2 hours at room temperature, washed in 1x PBS, then incubated in RIMS for 2 days at 4°C. The intestine was then mounted onto a slide in RIMS and imaged with the LSM 800 confocal microscope.

### Photoconversion

Adapting a previously described protocol (Hatta et al., 2006), we photoconverted all enteric neurons of Phox2b-kaede fish at 4.5 dpf using a Zeiss LSM 800 confocal microscope. Full thickness photoconversion was confirmed by post-conversion imaging through the full z-stack in all fish.

### Lipophilic dye neural tube fills

The far-red lipophilic dye DiIC_18_(5)-DS [ThermoFisher Scientific D12730] was prepared according to manufacturer’s instructions and injections were performed by adapting a previously described protocol (Gutzman and Sive, 2009). Briefly, 2.3 nL of dye was injected at approximately 30 hpf into the anterior neuropore using a glass capillary needle affixed to a microinjector [Nanoliter 2000, World Precision Instruments], Imaging at 6 dpf was performed with a Zeiss LSM 800 confocal microscope.

### Two-photon cell oblation

Adapting a previously described protocol (Muto and Kawakami, 2018), we ablated 10 enteric neurons within the distal hindgut (i.e. corresponding to the last two somite lengths of hindgut) of Pho2b-kaede fish at 4.5 dpf using a Zeiss LSM 710 confocal microscope with two-photon laser ablation.

### Drug exposure

Prucalopride [Millipore Sigma SML1371] was prepared at 10 uM and 100 uM in DMSO, and exposure occurred at 4.5 dpf through 5.5 dpf, unless otherwise stated. 4-OHT [Millipore Sigma H7904] was prepared at 20 uM in ethanol and exposure occurred at 3.5 dpf for a total of 16 hours. Controls in the prucalopride and 4-OHT experiments were exposed to equal volumes of DMSO or ethanol, respectively.

### Live-imaging

Live zebrafish larvae were anesthetized with tricaine and mounted within chamber slides using 1.2% low-melt agarose prepared in embryo water(“ZFIN: Zebrafish Book: General Methods,” n.d.). Additional embryo water was added after solidification of the agarose. All live-imaging was performed on a Zeiss LSM 800 confocal microscope with the incubator set at 28°C. For time-lapse experiments, z-stacks were collected every 4 minutes with a duration of 8 to 10 hours. Videos and 2D projections of Z-stacks were produced using Imaris software [Bitplane]. All other live-images were produced with ImageJ software.

For the functional assay, a continuous video was collected after placing the mounted larvae in the microscope’s heated incubator chamber for 30 minutes. Then, a baseline video was collected for 3 minutes in the z-plane corresponding to the mid-depth of the intestine, followed by addition of prucalopride in DMSO for a final concentration of 10 uM or DMSO alone to the individual fish’s chamber, and then a 15-minute video was immediately collected. An “expulsive contraction” was defined as a contraction resulting in the expulsion of autofluorescent intraluminal mucous out of the hindgut and into the external environment. Videos were produced using ImageJ software.

### Cell counting and statistics

Cell counting was performed manually using ImageJ software. In non-ablated fish, cell counts were within the hindgut corresponding to the last four somite lengths of hindgut. In the cell ablation experiments, cell counts were within the distal hindgut corresponding to the last two somite lengths of hindgut (i.e. within the field of ablation). Statistics were performed using Graphpad Prism 8 [Graphpad Software, Inc.] using Student t-test for 2 group comparisons and 1-way ANOVA for >2 group comparisons, with a P-value <0.05 indicating statistical significance.

## RESULTS

### Sox10-expressing cells and enteric glia are absent in the post-embryonic zebrafish intestine

*Sox10* is an early neural crest marker important for differentiation of nearly every neural crest lineage with the exception of cartilage, which instead uses its paralog *Sox9* (Martik and Bronner, 2017). *Sox10* marks early migratory neural crest cells and is retained after differentiation by enteric glia as well as melanocytes, but lost from enteric neurons (Heanue and Pachnis, 2007). During embryogenesis, vagal neural crest cells expressing *Sox10* delaminate from the neural tube, invade the foregut, and migrate in a generally rostral to caudal fashion along the intestine (Lake and Heuckeroth, 2013; Rao and Gershon, 2018). In chick and murine models, these enteric vagal neural crest cells proliferate as they migrate, and a proportion of the daughter cells cease migration and differentiate into neurons and glia. As vagal neural crest cells differentiate into enteric neurons, they downregulate *Sox10* expression and express *Phox2b* and other neuronal differentiation markers (Heanue and Pachnis, 2007); in contrast, enteric glia maintain *Sox10* expression and upregulate GFAP, S100B, and PLP1 (Gulbransen and Sharkey, 2012; Rao et al., 2015). In zebrafish, enteric vagal neural crest cells complete their colonization of the hindgut by 3 days post fertilization (dfp) (Heanue et al., 2016).

Previous studies have hypothesized that enteric neurogenesis is maintained postnatally, arising either from resident enteric neuronal precursors or enteric glia (Joseph et al., 2011; Kulkarni et al., 2017; Laranjeira et al., 2011). In search of such resident progenitors in the larval zebrafish intestine, we first performed ISH for *Sox10* and the enteric glial marker *PLP1a*. Surprisingly, while *Sox10* signal was identified as expected on migrating vagal neural crest cells within the intestine at embryonic stages, it was down-regulated in the intestines of 3.5 dpf larvae, corresponding with *Phox2b* expression throughout the gut. Furthermore, *PLP1a* transcripts also were absent in the intestine at both stages, albeit present in other parts of the nervous system [Fig. 1a-d].

**Figure 1:**
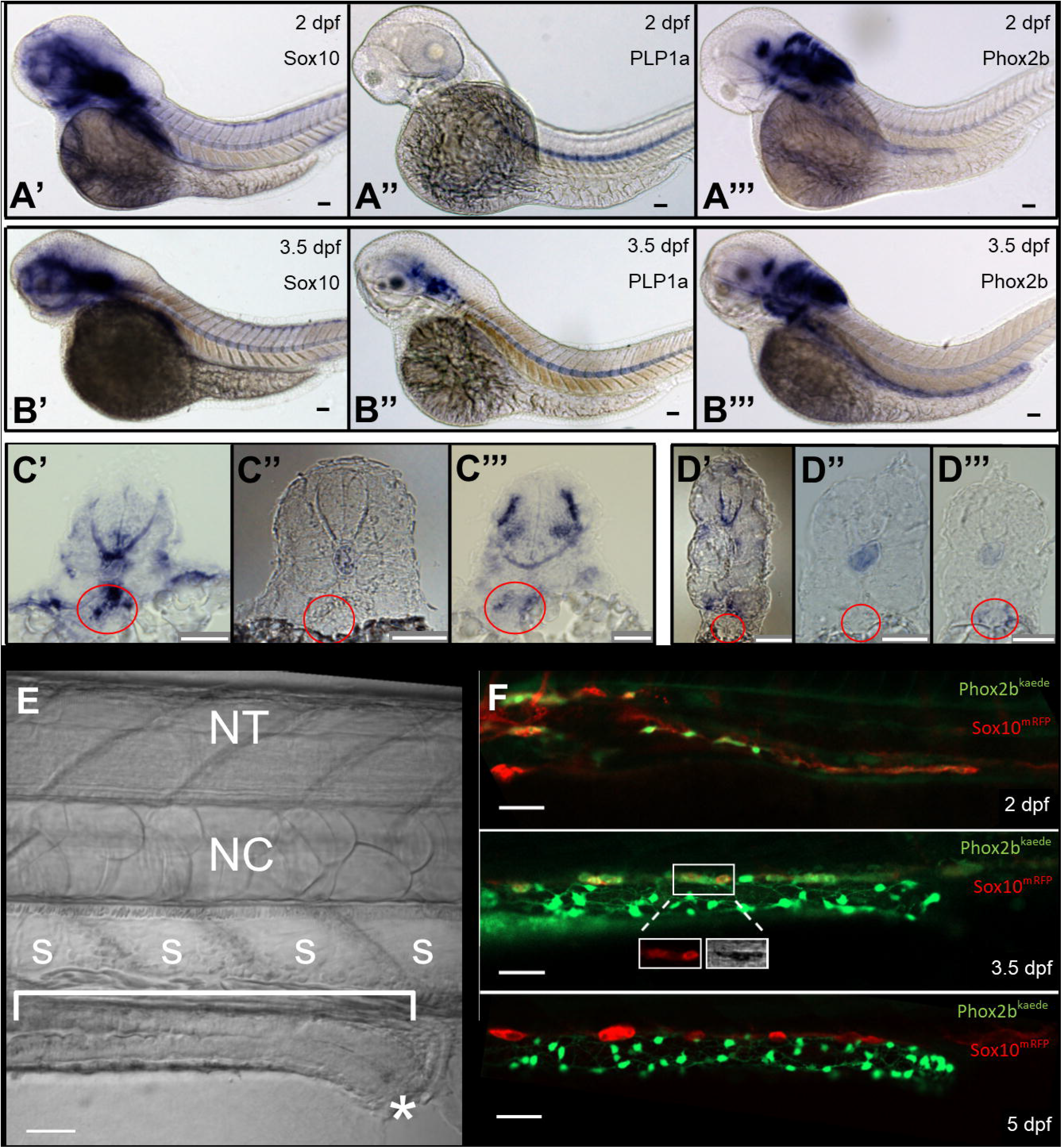
Resident neuronal progenitors are absent in the post-embryonic intestine. 1A) ISH of enteric neural elements in 2 dpf embryos. *Sox10* is detected at 2 dpf as a stream in the midgut that does not yet extend to the hindgut (A’), *PLP1a* exhibits no expression (A”), and *Phox2b* has weak expression in the proximal gut (A’“). Expected probe trapping is evident in the notochord. 1B) ISH of enteric neural elements in 3.5 dpf larvae. *Sox10* signal is absent in the intestine, though proximal probe trapping is seen in the nascent swim bladder (B’). *PLP1a* is expressed dorsally, but no expression is evident in the intestine (B”). *Phox2b* expression extends throughout the intestine (B’”). 1C) Proximal cross sections of 2 dpf embryos stained for Sox10 (C’), *PLP1a* (C”), and *Phox2b* (C’”), with the nascent foregut marked by a red circle. 2) Distal cross sections of 3.5 dpf larvae stained for Sox10 (D’), *PLP1a* (D”), and *Phox2b* (D’”), with the developing midgut marked by a red circle. 1E) Anatomic orientation; fluorescent figures are oriented in this manner unless otherwise stated. The intestine is located ventrally (bracket) and extends anterior (left) to posterior (right), ending at the anus (star). A row of polygonal somites (s) are arranged dorsal to the intestine. The notochord (NC) and neural tube (NT, not visible in this image) are located dorsally. 1F) Live imaging of Phox2b-kaede x Sox10-mRFP fish are consistent with ISH results: a migrating chain of Sox10 cells is observed in the midgut that does not yet extend to the hindgut at 2 dpf, but then Sox10 expression ceases at 3.5 and 5 dpf. A few Sox10-expressing cells are seen dorsolateral to the intestine are consistent with melanocytes, as supported by visible pigment in TPMT (inset, 3.5 dpf panel). Scale bars: 50 um

We next performed live-imaging using the transgenic line Sox10-mRFP (Kucenas et al., 2008) crossed with Phox2b-kaede (Harrison et al., 2014). As expected, Sox10 labelled cells were observed migrating along the intestine at 2 dpf, with only sparse Phox2b co-expression in the proximal foregut. In contrast, by 3.5 dpf, Sox10 was no longer expressed within the intestine (though Sox10-positive melanocytes were identified dorsolateral to the intestine). By 5 dpf, a conspicuous neuronal plexus expressing Phox2b-kaede had formed but no Sox10 expression was observed in the intestine [Fig. 1f, Supp.1]. Together, these results show that from 3.5 dpf onward, Sox10-expressing cells appear to be absent from the zebrafish intestine and confirm our ISH results, suggesting that there are no Sox10 expressing cells resident in the intestine at 3.5 dpf.

Next, we employed an indelible Cre transgenic line, Tg(Sox10:GAL4-UAS-Cre;ubi:switch), which permanently labels all Sox10-derived lineages with mCherry (Cavanaugh et al., 2015). Fish were euthanized and fixed at 5 dpf and then immunostained for the neuronal marker HuC/D and the Cre reporter mCherry. We found all Cre labelled cells co-localized with HuC/D, but no Cre-labelled cells were HuC/D-negative [Fig.2] indicating that 1) all Sox10-derived cells within the intestine have differentiated into enteric neurons by this stage, and 2) there are no non-neuronal Sox10-derived cells (i.e. resident precursors or glia) in the post-embryonic intestine. Of note, at this developmental stage, all Phox2b-kaede expressing cells co-localize with HuC/D [Supp.2], indicating that these cells are committed to a neuronal lineage.

**Figure 2:**
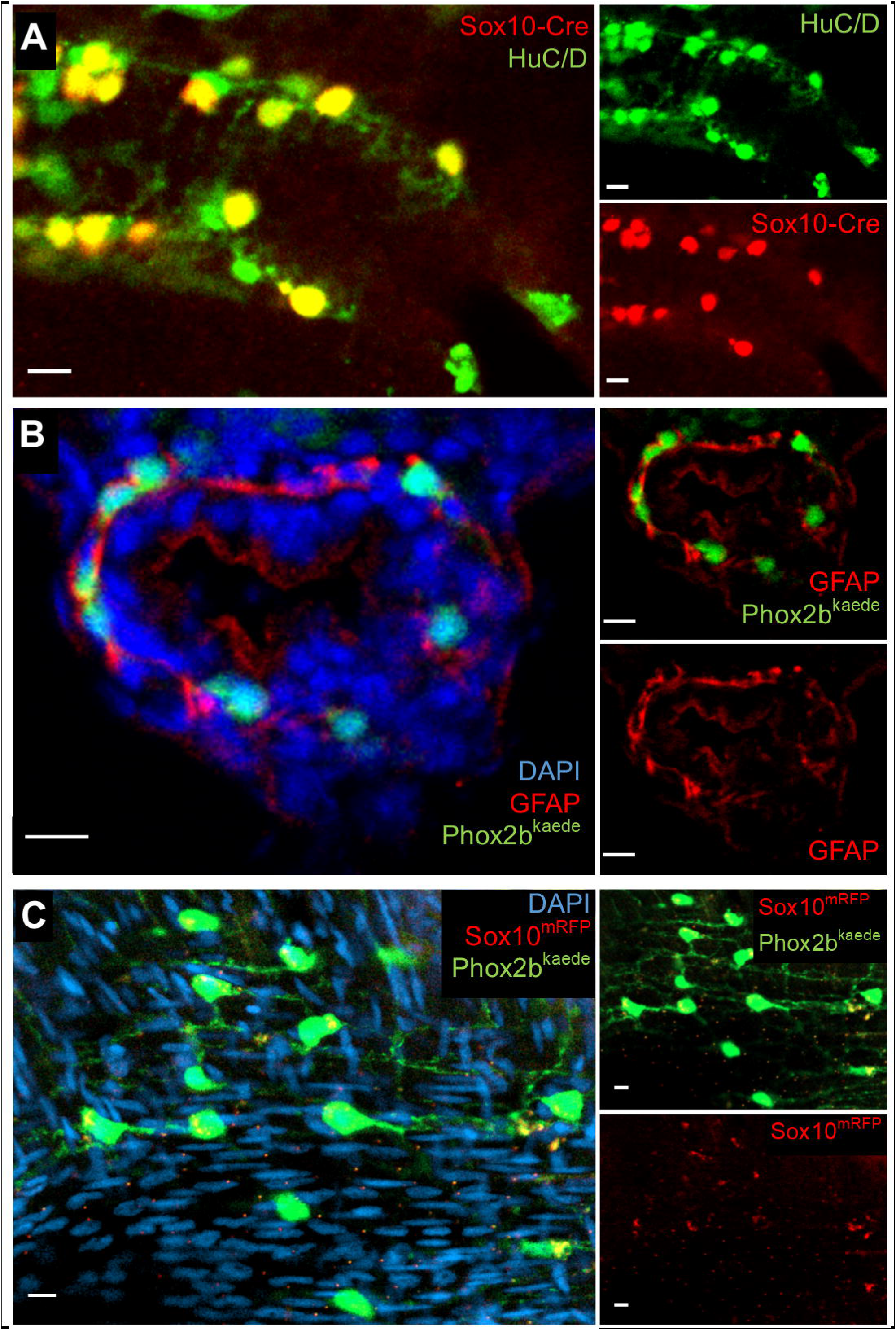
Further assays in larvae and adult support an absence of resident neuronal progenitors and enteric glia in the intestine. 2A) Lineage tracing with an indelible Sox10-Cre line suggests enteric neurons are the sole fate of enteric vagal neural crest cells. At 5 dpf, fish were fixed and underwent IHC for the Cre reporter, mCherry, and the neuronal marker HuC/D. All Cre-labelled cells co-localized with HuC/D, and no Cre-positive, HuC/D-negative cells were observed. 2B) IHC with GFAP does not demonstrate convincing enteric glial cell bodies. Phox2b-kaede fish were fixed at 5 dpf, and axially sectioned for IHC for GFAP, a glial marker with cytosolic expression. Imaging of the endogenous Phox2b-kaede signal in concert with the GFAP IHC revealed a fibrillary pattern of GFAP closely associating with enteric neurons and other cells, which likely represents projections from extrinsic glia. 2C) Whole-mount imaging of adult zebrafish intestine suggests that enteric glia and resident neuronal progenitors do not form later in development. Adult intestine from Phox2b-kaede x Sox10-mRFP fish that underwent optical clearing with RIMS revealed numerous enteric neurons, but no cell bodies expressing Sox10. Extrinsic glial projections are suggested by a fibrillary pattern of Sox10 expression. Scale bars: 10 um

An antibody to GFAP has previously been used as a marker to suggest the presence of enteric glia in zebrafish intestine. Therefore, we performed immunohistochemistry on 5 dpf Phox2b-kaede larvae sections using an antibody against zebrafish GFAP. As shown previously (Baker et al., 2019; Hagström and Olsson, 2010), we found GFAP immunoreactivity within the intestine; however, the GFAP appeared to be associated with cell processes but absent from cell bodies within the intestine [Fig.2b]. These findings likely reflect projections from extrinsic fibers but not resident cells within the intestine.

Lastly, to determine if enteric gliogenesis occurs later in development, we performed wholemount imaging of RIMS-cleared adult zebrafish intestine from the Sox10-mRFP x Phox2b-kaede line. While numerous Phox2b-kaede cells were present within the muscularis, no Sox10 cell bodies were identified, though RFP-positive signal corresponding to cell projections was observed [Fig.2c].

Taken together, these data suggest that vagal neural crest-derived cells within the zebrafish intestine all differentiate into neurons, with apparent absence of resident glia or progenitors.

### Enteric neurogenesis persists in the post-embryonic intestine in normal development and after injury

Given the continued growth of the intestine through adulthood, we hypothesized that enteric neurogenesis persists in post-embryonic stages. To test this, we employed the photoconvertible Phox2b-kaede line which upon exposure to light converts from green to red. We photoconverted all kaede-labelled cells within the 4.5 dpf intestine, after the vagal neural crest has completely colonized the intestine, such that all neurons that were initially in the green fluorescent conformation [Fig.3a] were converted to red [Fig.3b]. At 5 dpf, these fish were re-imaged.

**Figure 3:**
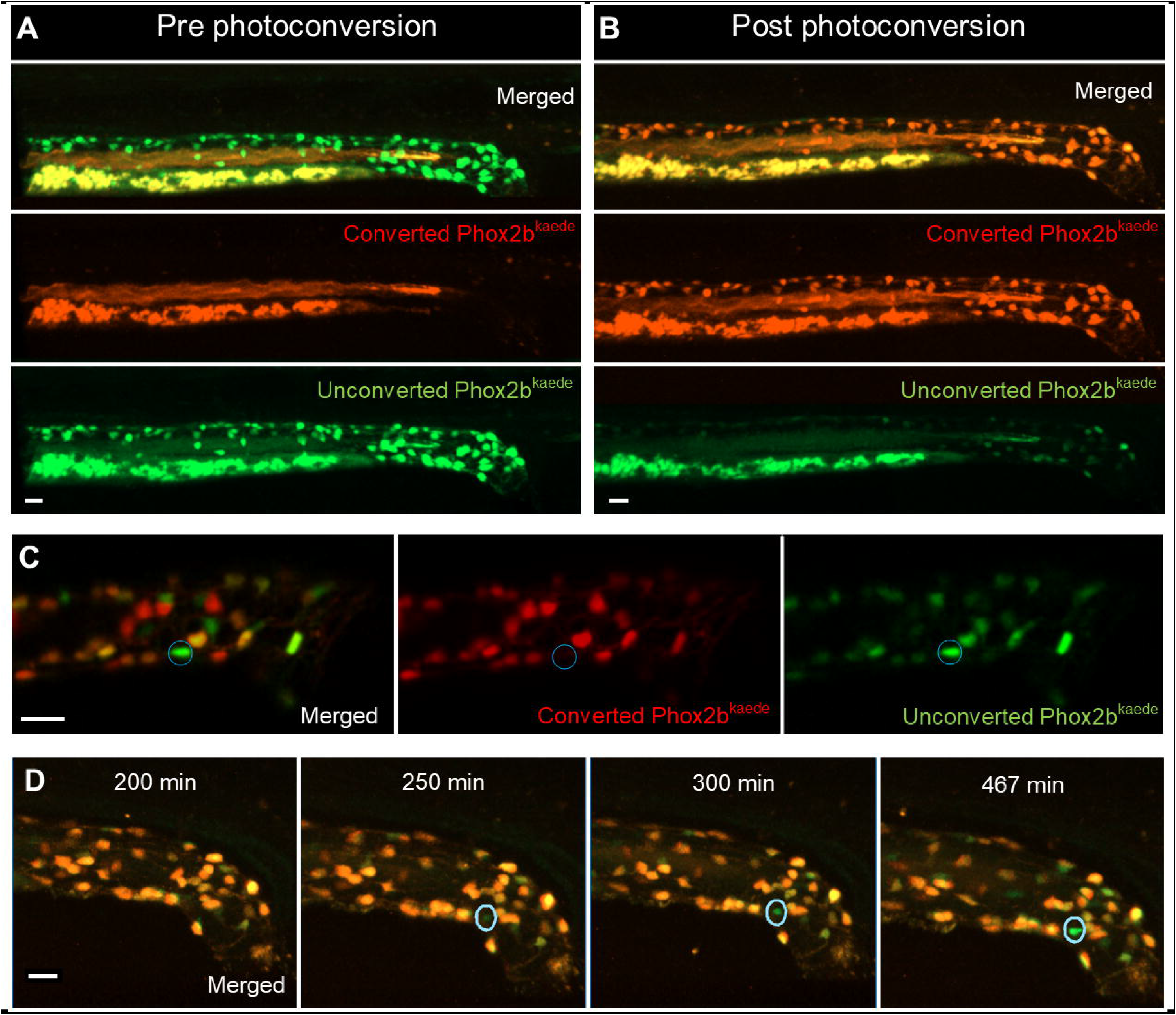
Enteric neurogenesis persists in the post-embryonic development despite an absence of resident neuronal precursors. 3A-B) 2D projection of z-stack from a 4.5 dpf Phox2b-kaede fish demonstrates green fluorescent enteric neurons, but no red fluorescent cells (3A). Yolk and intraluminal mucous exhibit expected autofluorescence in both channels. After photoconversion of all Phox2b-kaede neurons in the gut, all enteric neurons fluoresce red, though some retain decreased green fluorescence (3B). 3C) Live imaging 12 hours after photoconversion at 4.5 dpf reveals the appearance of green fluorescent enteric neurons in the intestine with no red fluorescence, indicating that these neurons did not arise from pre-existing red fluorescent Phox2b-kaede cells. 3D) Live 2D projection of a 10-hour time-lapse after photoconversion at 4.5 dpf detects the emergence of de novo enteric neurons, as indicated by the gradual appearance of a green-only neuron in a region of the intestine that was originally not occupied by a neuron. Scale bars: 20 um

Interestingly, we noted the appearance of Phox2b+ cells that only had green fluorescence [Fig.3c], suggesting they were newly born enteric neurons that did not arise from pre-existing Phox2b cells. To further validate this, we performed a 10 hour live time-lapse imaging experiment after photoconversion of all Phox2b-kaede cells and captured the emergence of de novo, green-only Phox2b-kaede enteric neurons [Fig.3d, Supp.3].

Next, we examined whether loss of existing enteric neurons was followed by neurogenesis. Using the Phox2b-kaede line, we conducted two-photon laser ablation of 10 Phox2b-kaede cells in the distal hindgut of 4.5 dpf zebrafish [Fig.4a-b], and immediately photoconverted the remaining cells as described above. Upon re-imaging at 5dpf, we again detected de novo enteric neurons [Fig.4c]. Time-lapse imaging over 8 hours in a 4.5 dpf Phox2b-kaede fish that underwent laser injury of 10 distal hindgut enteric neurons followed by photoconversion of all remaining enteric neurons revealed an injured neuron involuting and then being replaced by an emerging Phox2b-kaede de novo cell that appeared to actively migrate and extend projections to nearby neurons [Fig.4d, Supp.4].

**Figure 4:**
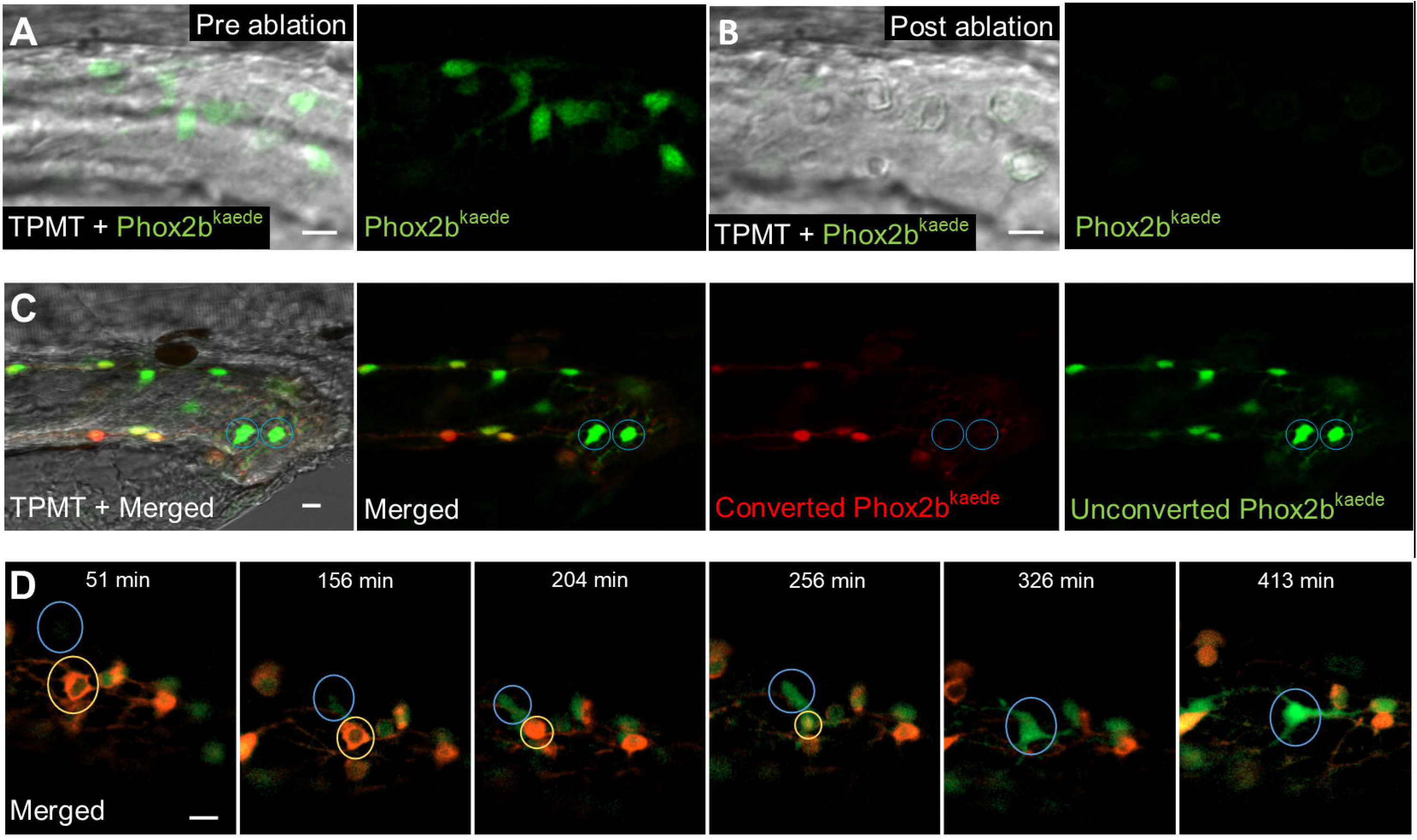
De novo enteric neurons replace ablated neurons in a post-embryonic injury model. 4A-B) Prior to 2-photon laser ablation, Phox2b-kaede enteric neurons are clearly visualized within the hindgut (A). After ablation, these neurons are no longer present in the hindgut, and TPMT reveals the injury site to be restricted to the neuron location (B). 4C) At 4.5 dpf, fish underwent laser ablation of 10 enteric neurons within the distal hindgut, followed by photoconversion of all remaining enteric neurons within the whole length of the gut. Live-imaging was performed 12 hours later, and detected multiple de novo, green fluorescent-only enteric neurons in the hindgut. 4D) 8-hour time-lapse of a fish at 4.5 dpf that underwent focal injury (but not complete ablation) of enteric neurons followed by pan-gut photoconversion reveals the involution of an injured neuron that is replaced by a de novo, green fluorescent-only enteric neuron. The new neuron initially appears very faintly at the dorsal-most aspect of the intestine but increases in intensity as it migrates to replace the involuted neuron and extends projections to neighboring neurons. Scale bars: 10 um

### Lineage tracing supports a trunk neural crest origin of post-embryonic enteric neurogenesis

Given that new Phox2b neurons apparently did not arise from existing neurons and there do not appear to be progenitors/glia in the intestine at this stage, we next investigated the possibility that these cells may arise from extrinsic sources. To explore the possibility that these come from the trunk spinal cord from which some Schwann cell precursors arise, we performed lineage tracing with the lipophilic dye DilCis(5)-DS, which fluoresces in the far-red wavelength. To this end, we injected dye into the neural tube of Phox2b-kaede embryos at approximately 30 hours post fertilization (hpf), after the vagal neural crest has completed emigration from the neural tube [Supp.5]. Live-imaging of injected fish at 6 dpf revealed numerous dye-labelled enteric neurons: of 30 Dil-injected fish, 15 had Phox2b-kaede enteric neurons that co-localized with the dye (mean: 4.60 dye-labelled enteric neurons per fish, SD: 2.29) [Fig.5a-b]. There was no statistically significant difference in the distribution of dye-labelled enteric neurons within the foregut, midgut, or hindgut. Given that trunk neural crest cells migrate from the neural tube during this time frame, these findings suggest that trunk, but not vagal, neural crest-derived cells are the source of these new enteric neurons.

**Figure 5:**
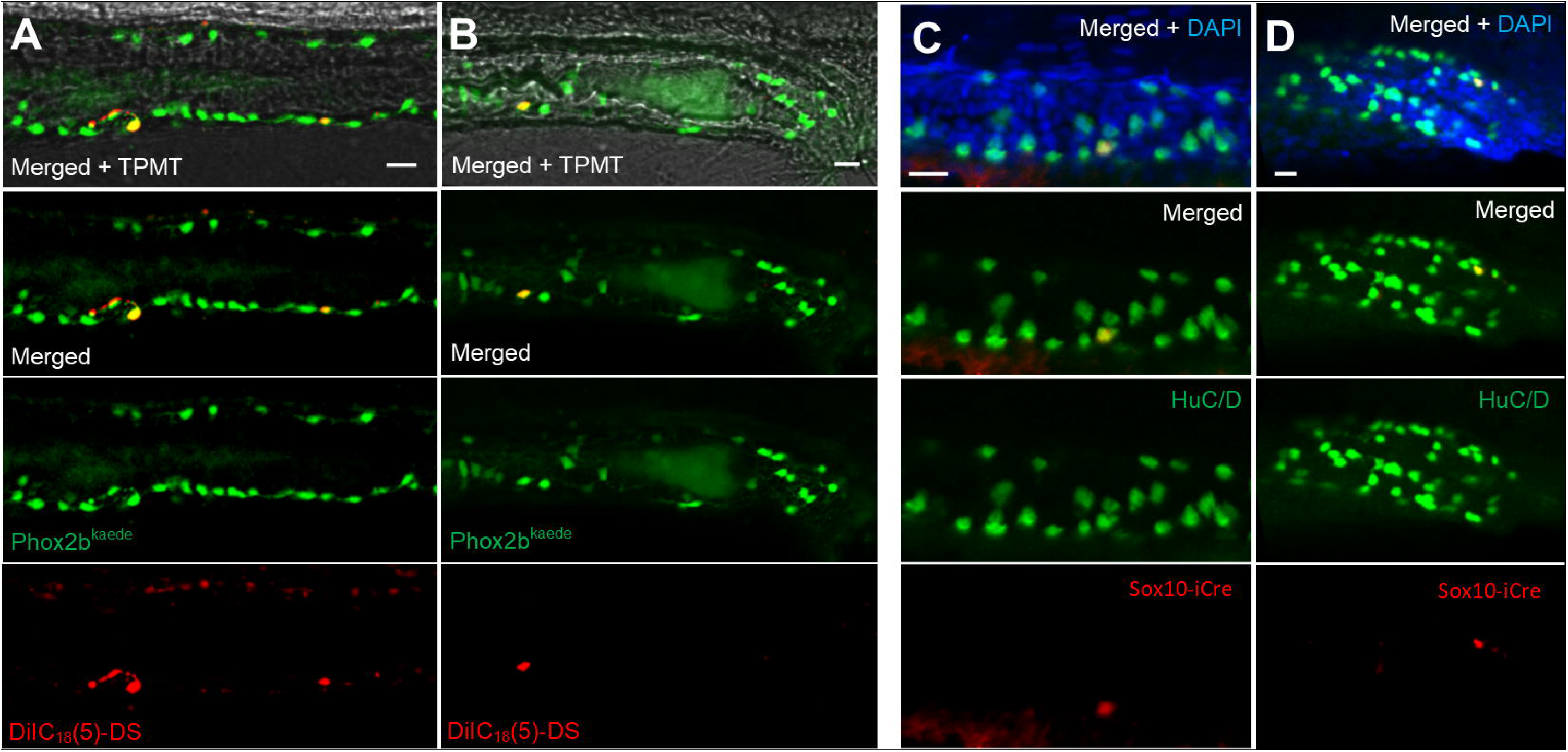
Lineage tracing demonstrates a trunk neural crest origin of post-embryonic neurogenesis. 5A-B) Phox2b-kaede embryos underwent neural tube injections of a far-red lipophilic dye at 30 hpf, after vagal crest has delaminated from the neural tube. Live images at 6 dpf of the midgut (5A) and hindgut (5B) demonstrate Phox2b-kaede enteric neurons that co-localize with the dye, indicating their trunk origin. 5C-D) Fish from the inducible Sox10-Cre line were exposed to 4-OHT at 3.5 dpf and underwent IHC for the Cre reporter, mCherry, and the neuronal marker, HuC/D at 5.5 dpf, with Cre labelled enteric neurons observed in the midgut (5C) and hindgut (5D). As Cre induction occurred after Sox10 is no longer present within the intestine, these results support a trunk neural crest origin of these enteric neurons. Scale bars: 20 um

Next, we performed lineage tracing using transgenic approaches with the inducible Sox10-Cre line, Sox10ERT2 x ubi:switch (Mongera et al., 2013). Cells expressing Sox10 during the induction period are permanently labelled with the reporter, mCherry. Zebrafish were induced at 3.5 dpf (after vagal crest has completed intestinal colonization and Sox10 expression is no longer observed in the intestine) for a total of 16 hours and then fixed at 5.5 dpf [Supp.6]. These fish then underwent immunostaining using neuronal marker HuC/D and the Cre reporter, mCherry. While this line only labels a subset of Sox10 expressing cells, the results revealed enteric neurons that co-localized with the Cre reporter [Fig.5c-d]. Of the 34 induced fish, 11 exhibited Cre-labelled enteric neurons (mean: 1.63 Cre-labelled enteric neurons per fish, SD: 0.67). These results confirm that these neurons arose from a neural crest-derived source external to the intestine.

Taken together, our two lineage tracing experiments provide evidence that de novo enteric neurogenesis arises from trunk neural crest derived progenitors, likely to be Schwann cell precursors that originate from the trunk neural tube and migrate to the intestine.

### 5HT4R agonism promotes post-embryonic enteric neurogenesis

Previous studies in rodents (Liu et al., 2009; Matsuyoshi et al., 2010) have shown that postnatal enteric neurogenesis occurs after exposure to 5HT4R agonists. Recently, the highly specific 5HT4R agonist prucalopride has been approved for clinical use in the United States (“Drug Approval Package,” n.d.) to treat slow transit constipation as this drug stimulates pro-motility activity of enteric neurons (Wong et al., 2010).

Consistent with this, using zebrafish transgenic line HuC:H2B-GCaMP6 (Freeman et al., 2014), we found that fish exposed to 10 uM prucalopride exhibited significantly increased hindgut contractions resulting in increased intraluminal expulsion at 5 dpf compared to controls (mean: 4 vs 0.5 expulsive contractions; p <0.001) [Fig.6a-b, Supp.7-8]. Notably, these contractions appeared to be associated with increased GCaMP activity in enteric neurons, suggesting neuronally-mediated contractions. The results from this functional assay demonstrate that 5HT4R signaling in zebrafish is active at these drug concentrations.

**Figure 6:**
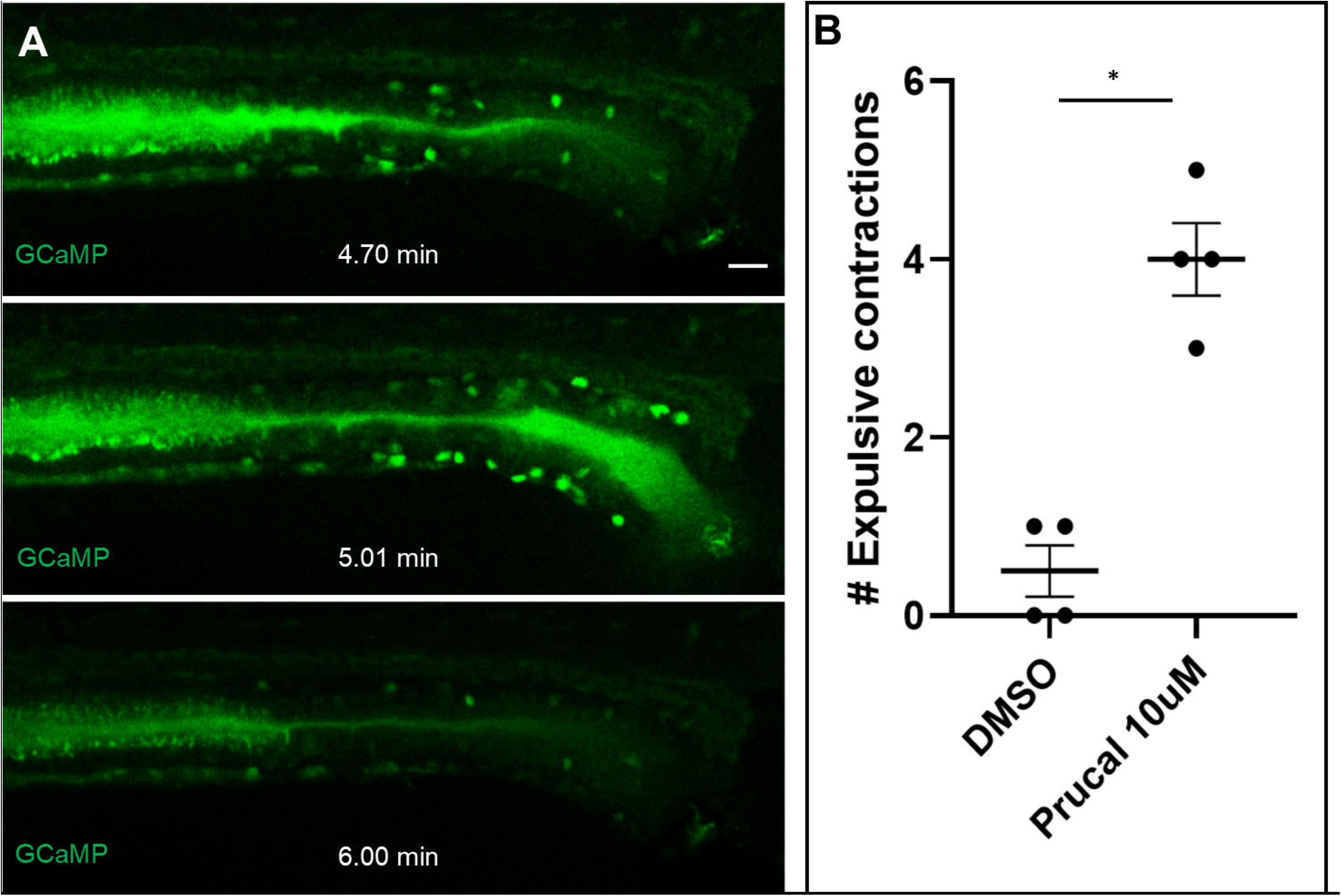
Prucalopride is active in the zebrafish and increases intestinal motility. 6A) Stills from a video of a live HuC-H2B GCaMP6 fish exposed to 10 uM prucalopride at 5 dpf reveals increased intestinal motility, as measured by expulsive contractions of autofluorescent intraluminal mucous into the external environment (0.5 vs 4.0; p=0.0002). Increased GCaMP signal was observed in association with expulsive contractions, suggesting neuronally-mediated motility. 6B) Compared to controls, fish exposed to prucalopride exhibited significantly more expulsive contractions. Scale bar: 20 um

To assess the effects of prucalopride on post-embryonic enteric neurogenesis, we utilized the Phox2b-kaede line and photoconverted all enteric neurons at 4.5 dpf. Cohorts of these fish were then exposed to 10 uM prucalopride, 100uM prucalopride, or DMSO [Supp.9A], Live-imaging was performed at 5 dpf. The results show that both prucalopride treated cohorts possessed significantly more de novo enteric neurons (green-only) in the hindgut compared to controls (control: 14, 10 uM: 25, 100 uM: 27.67; p<0.05) [Fig.7a]. As there was no difference in de novo enteric neuron numbers between the 10 uM and 100 uM cohorts, 10 uM of prucalopride was the dose used in subsequent experiments.

**Figure 7:**
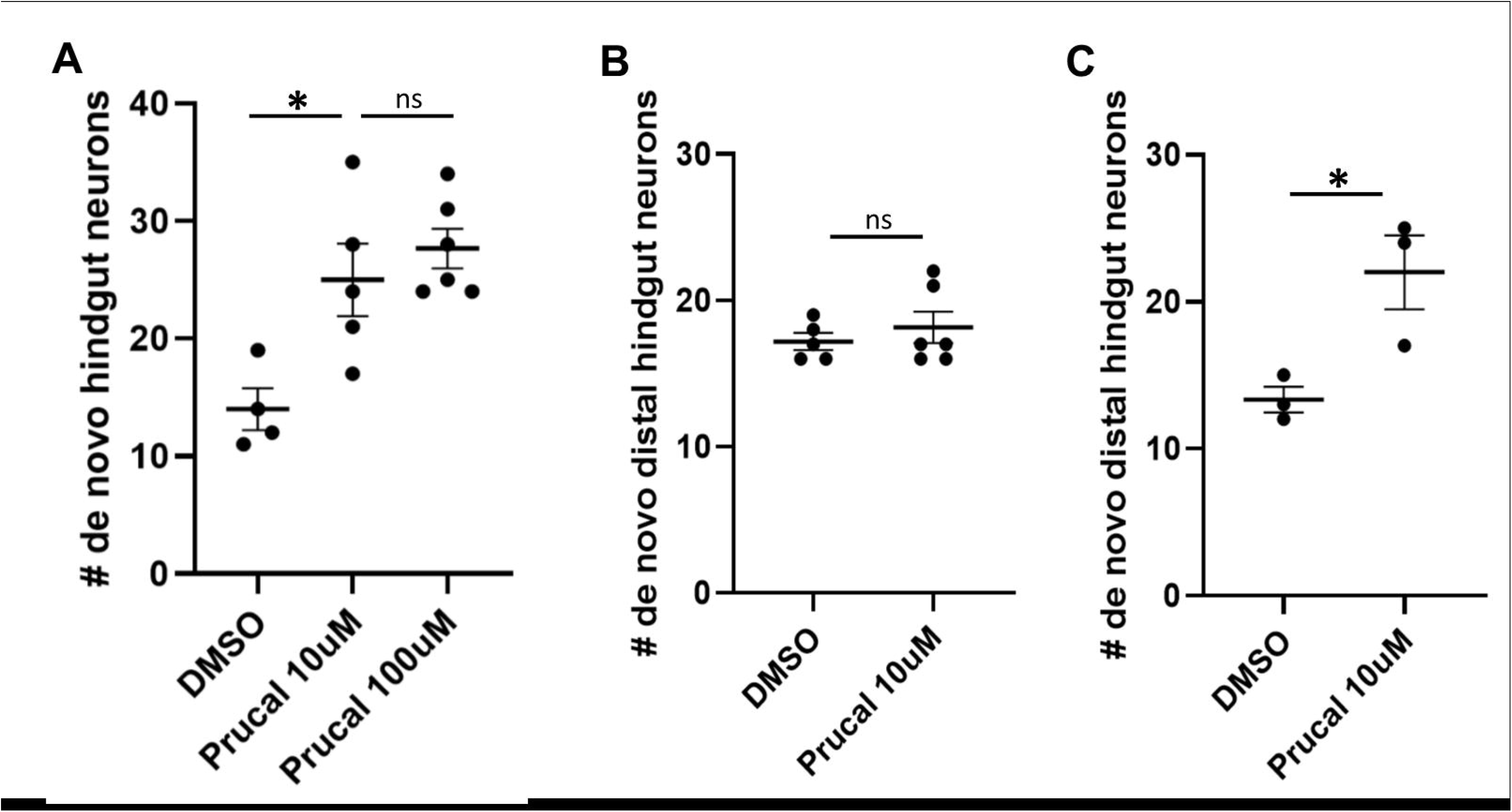
Prucalopride promotes enteric neurogenesis in normal development and injury. 7A) After photoconversion of all enteric neurons at 4.5 dpf, cohorts of Phox2b-kaede fish were exposed to 10 uM prucalopride (N=5), 100 uM prucalopride (N=6), or DMSO (N=4) for 12 hours and then live-imaged at 5 dpf. The mean number of de novo hindgut neurons was significantly higher in fish treated with prucalopride (25, 27.7, and 14, respectively; p=0.019 and p=0.0036, respectively). 7B) At 4.5 dpf, Phox2b-kaede fish underwent laser ablation of 10 distal hindgut enteric neurons and then photoconversion of all enteric neurons. Cohorts were exposed to 10 uM prucalopride (N=6) or DMSO (N=5) for 12 hours and then live-imaged at 5 dpf. There was no difference in de novo distal hindgut neurons between these two groups. 7C) Under similar experimental design as 7C, fish were instead exposed to prucalopride 10 uM (N=3) or DMSO (N=3) at 3.5 dpf for 12 hours, and then underwent cell ablation and photoconversion at 4.5 dpf. At 5 dpf, live-imaging revealed significantly more distal hindgut neurons in fish pre-treated with prucalopride (22 vs 13.3; p=0.031).

To assess if 5HT4R agonism is involved in enteric neurogenesis after injury, we performed two-photon laser ablation of enteric neurons in the hindgut of Phox2b-kaede fish at 4.5 dpf followed by photoconversion of all enteric neurons within the intestine. Subsequently, one cohort of these fish were treated with 10 uM prucalopride and a control group was treated with DMSO [Supp.9B], Fish were reimaged at 5 dpf, and de novo enteric neurons within the distal hindgut were counted; no significant difference was noted between the two groups (17.2 vs 18.2; p=0.47) [Fig.7b].

We next repeated this experiment but treated one cohort of fish with 10 uM prucalopride for 24 hours *prior to* laser ablation and photoconversion at 4.5 dpf [Supp.9C]. Compared to controls pre-treated with DMSO, pre-treatment with prucalopride resulted in significantly more (13.3 vs 22; p=0.03) de novo enteric neurons in the distal hindgut after injury [Fig.7c].

These findings suggest that exposure to prucalopride prior to injury promotes regeneration of enteric neurons, whereas a short course of treatment after injury has no effect on neurogenesis.

## DISCUSSION

In this study, we provide evidence that zebrafish enteric neurogenesis persists in the post-embryonic intestine both during normal development and after ablation of enteric neurons despite an apparent absence of enteric glia and/or Sox10-derived resident progenitors. Rather, lineage tracing experiments support the intriguing possibility that trunk crest-derived neural crest stem cells, likely to be Schwann cell precursors that migrate along nerves from the spinal cord to the intestines, are a source of this post-embryonic enteric neurogenesis. Along with the expected pro-motility effect, we also demonstrated that 5HT4R agonism with prucalopride increased post-embryonic neurogenesis in normal development and appeared to promote regeneration of enteric neurons if the exposure occurred prior to injury.

Our results are consistent with studies in the basal jawless vertebrate lamprey (Green et al., 2017) and mice (Uesaka et al., 2015), showing that there is an important contribution of Schwann cell precursors to the ENS during development. Moreover, our results suggest that this source persists post-embryonically to support ongoing neurogenesis reflecting both normal turnover of neurons and regeneration after injury. As lamprey lack a vagal neural crest, enteric neurogenesis from SCPs is likely to have been the basal state for populating the ancestral vertebrate intestine with neurons (Green et al., 2017). With the advent of the vagal neural crest in jawed vertebrates, they became the main embryonic source to populate enteric neurons whereas SCPs may have been repurposed as a means to supplement additional enteric neurons to accommodate continued growth during post-embryonic development as well as regeneration after injury.

The simplified zebrafish ENS compared with amniotes together with the apparent absence of enteric glia makes it a highly tractable model in which to examine the complex nature of post-embryonic enteric neurogenesis. Importantly, the zebrafish ENS develops in a homologous manner to humans during embryogenesis (Ganz, 2018; Heanue et al., 2016), and the intestine is anatomically and functionally segmented similarly to human small intestine and colon (Wang et al., 2010). Of note, the 5-HT_4_ receptor arose early in evolution (Hashiguchi and Nishida, 2007; Tierney, 2018), and 82% of disease-related human genes have a zebrafish homologue (Howe et al., 2013); thus, the zebrafish is an ideal system in which to explore fundamental features of post-embryonic enteric neurogenesis.

As SCP-derived enteric neurogenesis is conserved in mammals, our demonstration that this source of enteric neurogenesis may be amenable to pharmacologic manipulation deepens the rationale for further exploration into 5HT4R-based therapies for human enteric neuropathies. Other 5HT4R agonists such as mosapride (not available in the United States) and tegaserod (limited use in the United States due to off-target side effects) have previously supported the role of this signaling pathway in enteric neurogenesis (Liu et al., 2009; Matsuyoshi et al., 2010). Our study is the first to employ prucalopride, a highly specific 5HT4R agonist that has recently been approved for use in the United States (“Drug Approval Package,” n.d.; Wong et al., 2010). Prior studies suggested 5HT4R agonism mediated its enteric neurogenic effect through a resident progenitor, but the evidence was inconclusive as a gut-extrinsic source was not assessed (Goto et al., 2013; Katsui et al., 2009; Liu et al., 2009; Matsuyoshi et al., 2010). Indeed, in one of these studies (Liu et al., 2009), neuronal precursors were first detected outside of enteric ganglia and then appeared to migrate within the ganglia, which has lead us and others (Uesaka et al., 2015) to hypothesize that these observations are consistent with SCP-derived enteric neurogenesis. While we found pre-treatment with prucalopride promoted enteric neurogenesis after injury, treatment after injury did not. This may suggest a period of recovery or a longer treatment duration is required to promote enteric neuronal regeneration, but further investigation is required.

There is conflicting evidence in the literature regarding the presence of enteric glia in zebrafish. Some authors have reported GFAP immunoreactivity in the intestine (Baker et al., 2019; Doodnath et al., 2010; Hagström and Olsson, 2010), leading them to conclude the presence of enteric glia. However, the observed immunoreactivity is fibrillary and likely to reflect projections from extrinsic glia or other cells types, as no cell bodies are evident. Furthermore, S100β, an enteric glial marker with nuclear expression, failed to reveal enteric glia in the zebrafish (Germanà et al., 2008). However, we cannot exclude the possibility of a vagal crest-derived Sox10-negative resident enteric neuronal progenitor, as suggested by one study (Kulkarni et al., 2017). On the other hand, our lineage tracing experiments using an indelible Sox10-Cre line, dye-labeling, and an inducible Sox10-Cre line support a gut-extrinsic, trunk neural crest-derived source of enteric neurogenesis.

Other studies have raised questions about the functional importance of glia in the mammalian ENS. For example, in male mice from which all enteric glia were genetically ablated, no obvious differences were observed in intestinal motility or predilection to enterocolitis (Rao et al., 2017). As zebrafish have a functional intestine despite the apparent absence of enteric glia, when and why enteric glia evolved represents an interesting question for future study. Considering that humans likely possess hundreds of millions of enteric glia (Grubišić and Gulbransen, 2017), clarifying their functional significance carries broad implications in gastroenterology (Gulbransen and Sharkey, 2012) and may be aided by investigating their evolutionary development.

Our simplified motility assay to assess prucalopride’s effect on the zebrafish intestine is easily accessible without the need for custom cameras or imaging processing programs, contrasting with other gastrointestinal motility studies in zebrafish (Ganz et al., 2018; Shi et al., 2014). The results demonstrate a functionally significant assay that is intuitively translatable to clinical end-points.

By applying live-imaging techniques in the zebrafish, we are able to specifically ablate individual enteric neurons with minimal damage to adjacent cells and then capture regeneration of enteric neurons in real time. Prior studies (Goto et al., 2013; Katsui et al., 2009; Laranjeira et al., 2011; Matsuyoshi et al., 2010) required the broad application of cytotoxic compounds or full-thickness surgical transection, followed by assessment for neurogenesis at later time points. Our approach has the advantage of specifically injuring enteric neurons followed by in vivo time-lapse detection of neurogenesis. Taken together, our results reveal fundamental features of post-embryonic ENS development and regeneration in the zebrafish, pointing toward potential therapeutic strategies to promote Schwann cell precursor-derived enteric neurogenesis in the treatment of enteric neuropathies.

## Abbreviations

ENS: enteric nervous system
SCP: Schwann cell precursor
ISH: in situ hybridization
IHC: immunohistochemistry
dpf: days post fertilization
hpf: hours post fertilization
5HT4R: 5-HT_4_ receptor

## Author contributions

WN El-Nachef: study concept and design; acquisition of data; analysis and interpretation of data; drafting of the manuscript; statistical analysis

ME Bronner: study concept and design; analysis and interpretation of data; critical revision of the manuscript; obtained funding; study supervision.

## ACKNOWLEDGEMENTS

We thank the Beckman Institute Biological Imaging Facility of Caltech for technical assistance with microscopy experiments. Support was provided by a UCLA Division of Digestive Diseases Seed Grant, the UCLASpecialty Training and Advanced Research Program, and NIH R35NS111564 and NIH R01NS108500. For generously supplying transgenic fish lines, we thank the Ian Shepherd Lab (Phox2b-kaede), the Jua-Nian Chen Lab (Sox10:GAL4-UAS-Cre and ubi:switch), and the Christiane Nüsslein-Volhard Lab (cmlc:GFP-Sox10:ERT2-Cre). Special thanks to Megan Martik and Can Li for sharing ISH probes (Sox10, Phox2bb). We also recognize Claire Hu (Caltech) for assistance with IHC.

## Notes

### Competing Interest Statement

The authors have declared no competing interest.

