## Supplemental figures and legends for "De novo enteric neurogenesis in post-embryonic zebrafish from Schwann cell precursors rather than resident cell types"

***[Video file attached separately]***

Supplement 1: Sox10 expression at 5dpf likely corresponds to melanocytes

A 2D projection of a z-stack collected at 5 dpf of a Phox2b-kaede x Sox10-mRFP fish hindgut reveals a linearly arrange collection of Sox10-expressing cells. However, 3-dimensional assessment indicates that these cells are located dorsolateral to the intestine and likely correspond to melanocytes which reside in this location.

Scale bar: 30 um

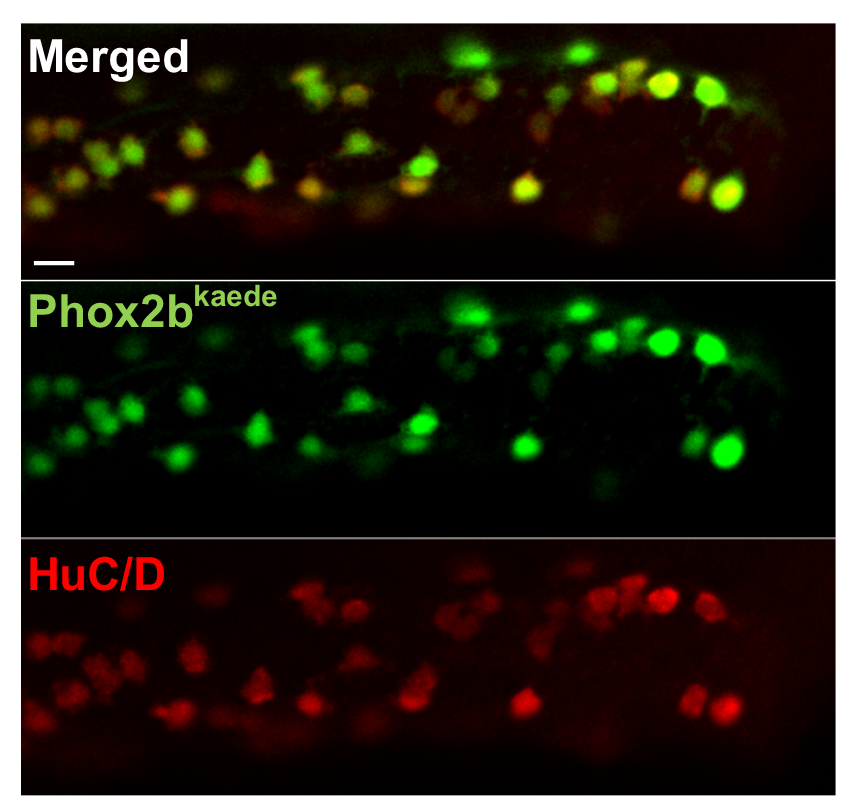

Supplement 2: HuC/D co-localizes with all Phox2b-kaede enteric neurons at 5 dpf.

Phox2b-kaede fish were fixed at 5 dpf and underwent IHC for HuC/D, and imaged for the endogenous kaede fluorescence and HuC/D. All Phox2b-kaede cells co-localized with HuC/D, indicating that at this stage, Phox2b represents differentiated enteric neurons.

Scale bar: 10 um

***[Video file attached separately]***

Supplement 3:

Video of the live time-lapse experiment from Figure 3D depicts the gradual appearance of a de novo enteric neuron in a portion of the hindgut that initially did not contain an enteric neuron. The de novo enteric neuron appears to make contact with neighboring cells.

Scale bar: 20 um

***[Video file attached separately]***

Supplement 4:

Video of the live time-lapse experiment from Figure 4D depicts the appearance of a de novo enteric neuron that initially appears very faintly in the dorsal periphery of the intestine. As an injured enteric neuron involutes, it is replaced by the migrating de novo enteric neuron, which gradually increases Phox2b-kaede expression and extends projections to nearby enteric neurons.

Scale bar: 10 um

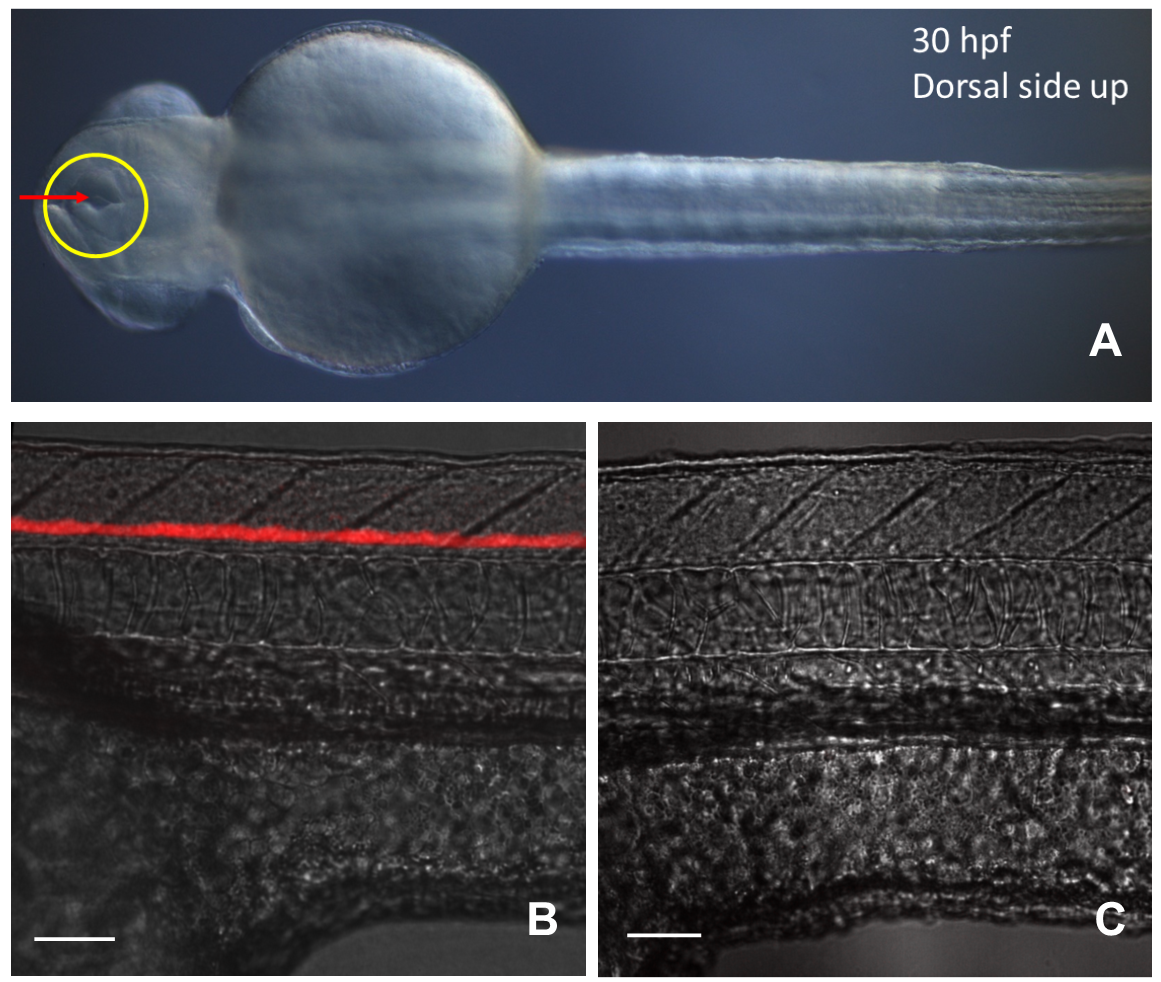

Supplement 5:

S5A) Schematic of lipophilic dye injection at 30 hpf. The anterior neuropore (yellow circle) is open at this time point, allowing insertion of a dye-filled capillary in the trajectory depicted by the arrow.

S5B-C) 1-hour post injection, a dye-colored stripe is present indicating successful neural tube fill. Control fish that were not injected did not exhibit far-red fluorescence.

Scale bar: 50 um

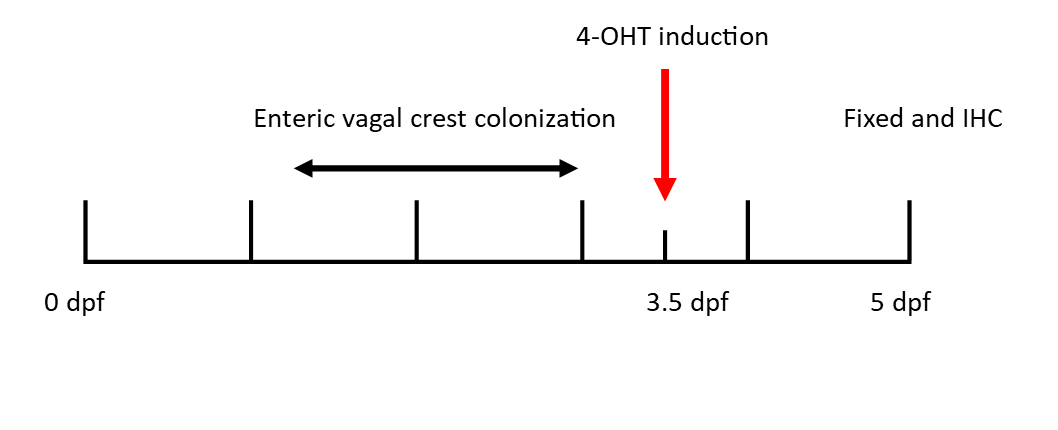

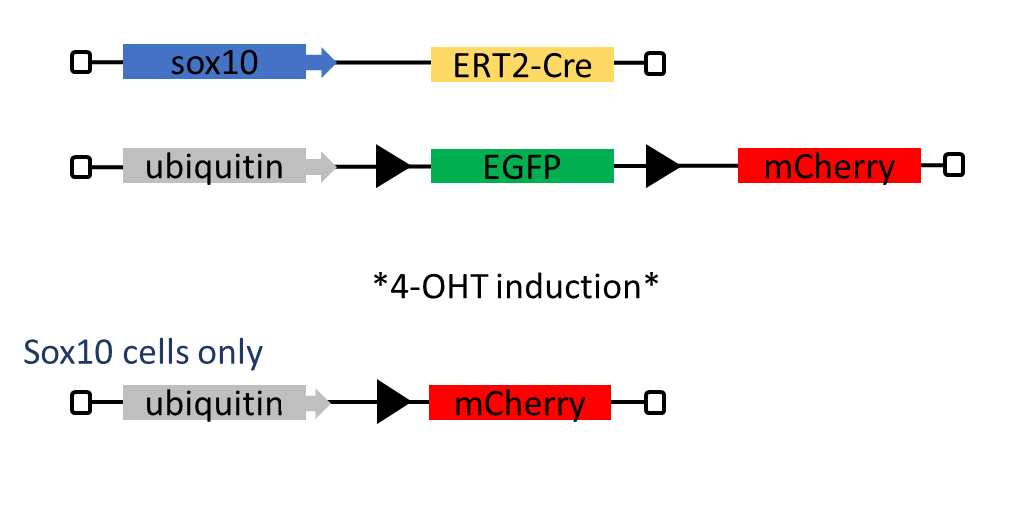

**A**

**B**

Supplement 6:

S6A) The inducible Sox10-Cre line was crossed with the reporter, ubi:switch. After exposure to the induction agent, 4-OHT, Cre is activated, cleaves the loxP sites specifically in cells expressing Sox10 at the time of induction, leading to those cells permanently being labelled by mCherry.

S6B) Schematic of the induction protocol for Figure 5C-D. Fish were exposed to 4-OHT at 3.5 dpf for 16 hours, when Sox10 is no longer expressed in the intestine. Cells labelled during this time are thus gut-extrinsic Sox10-expressing cells.

***[Video files attached separately]***

Supplements 7 and 8:

Video of a 5 dpf HuC-H2B GCaMP6 fish exposed to DMSO reveals low baseline motility over a 15 min time frame (Supp.7). In contrast, multiple expulsive contractions are observed with exposure to 10 uM prucalopride (Supp.8).

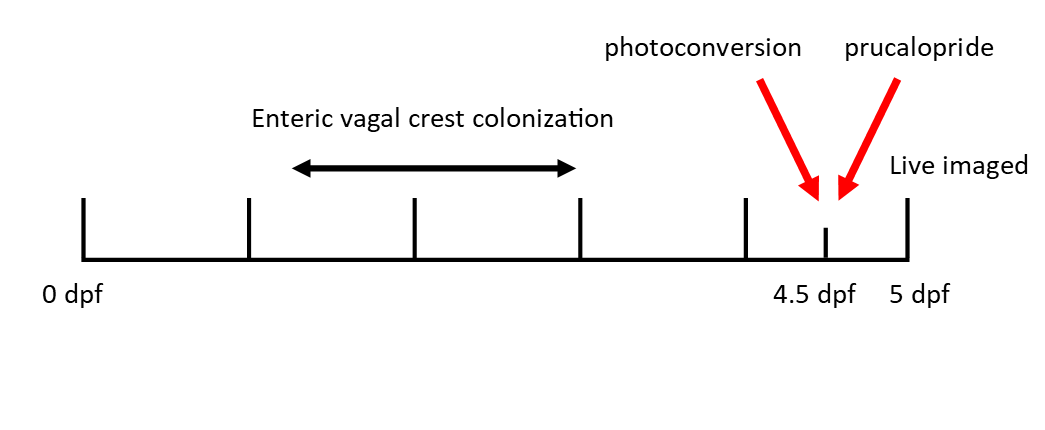

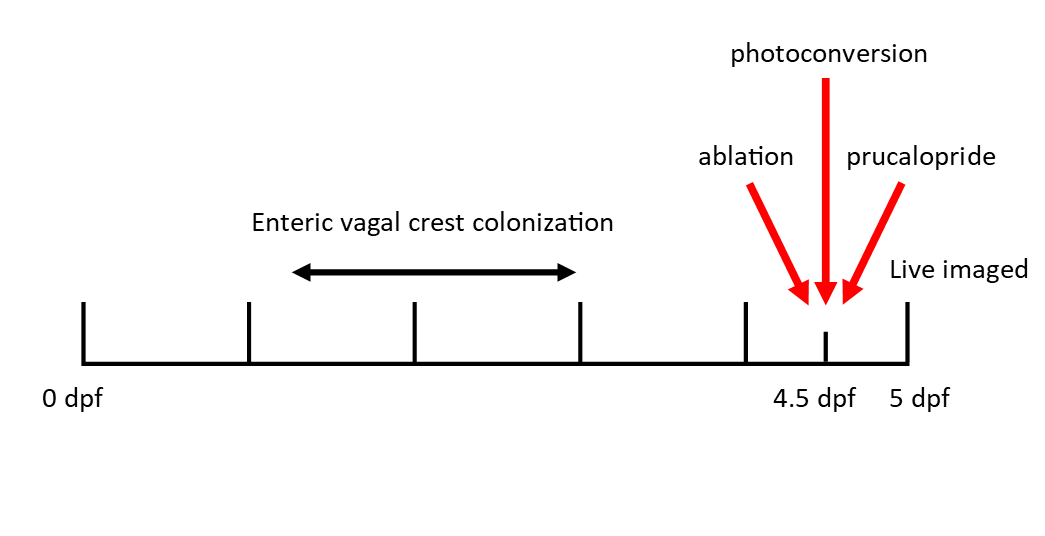

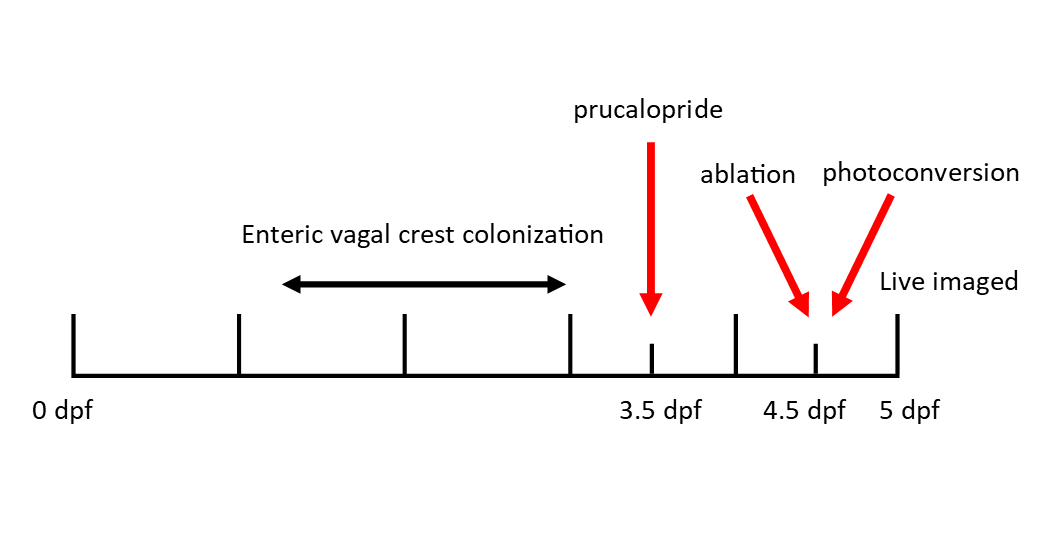

**A**

**C**

**B**

Supplement 9:

Supp.9A-C: Schematic of the protocols for Figures 7A, B, and C depicting the timing of drug exposure, photoconversion, and laser ablation.
